# Relationships between clans and genetic kin explain cultural similarities over vast distances: the case of Yakutia

**DOI:** 10.1101/168658

## Abstract

Archaeological studies sample ancient human populations one site at a time, often limited to a fraction of the regions and periods occupied by a given group. While this bias is known and discussed in the literature, few model populations span areas as large and unforgiving as the Yakuts of Eastern Siberia. We systematically surveyed 31,000 square kilometres in the Sakha Republic (Yakutia) and completed the archaeological study of 174 frozen graves, assembled between the 15^th^ and the 19^th^ century. We analysed genetic data (autosomal genotypes, Y-chromosome haplotypes and mitochondrial haplotypes) for all ancient subjects and confronted these to data on 190 modern subjects from the same area and the same population. Ancient familial links were identified between graves up to 1500 km apart, as well as paternal clans. We provide new insights on the origins of the contemporary Yakut population and demonstrate that cultural similarities in the past were linked to (i) the expansion of specific paternal clans, (ii) preferential marriage among the elites and (iii) funeral choices that could constitute a bias in any ancient population study.

## Introduction

Various mechanisms exist to explain how cultures spread among and between human populations (Alexrold 1997). In ancient populations however, it has proved a challenge to determine whether individuals themselves had moved or fashions had been transmitted between distinct groups (Guilaine and Crubézy, 2003). Palaeogenetics provide valuable additional information in such cases (Fernandez *et al.* 2014). We endeavoured to study clan and family relationships through genetics, to understand the expansion of Yakut culture in North-Eastern Siberia between the 17^th^ and 20^th^ centuries, from a few settlements pockets to more than 3 million km^2^ (Crubézy and Nikolaeva, 2017). During the last 15 years, we systematically surveyed 31,000 km^2^ (Figure 1) of especially suitable terrain in the Sakha Republic (Yakutia) and completed the archaeological study of 174 frozen graves assembled between the 15^th^ century and the 19^th^ century (Supplementary Data I). We categorised and analysed cultural characteristics and extracted genetic data (autosomal genotypes, Y-chromosome haplotypes and mitochondrial haplotypes) for 150 ancient subjects, along with modern data collected from 190 modern individuals from the same regions and the same populations. We organised the dataset according to all available criteria separately and then concurrently, in order to identify potential subsets within populations and specific links and differences between subjects. First, subjects were organised into cultural clusters based on the characteristics of the graves, then male lineage diversity was evaluated and compared to temporal and spatial information, finally close and distant genetic kinships were identified and studied in relation to geography and archaeology. These combined methods allow us to describe mechanisms of cultural transmission, interregional links through male-dependent clans and families, matrimonial preferences and burial choices in the ancient Yakut population.

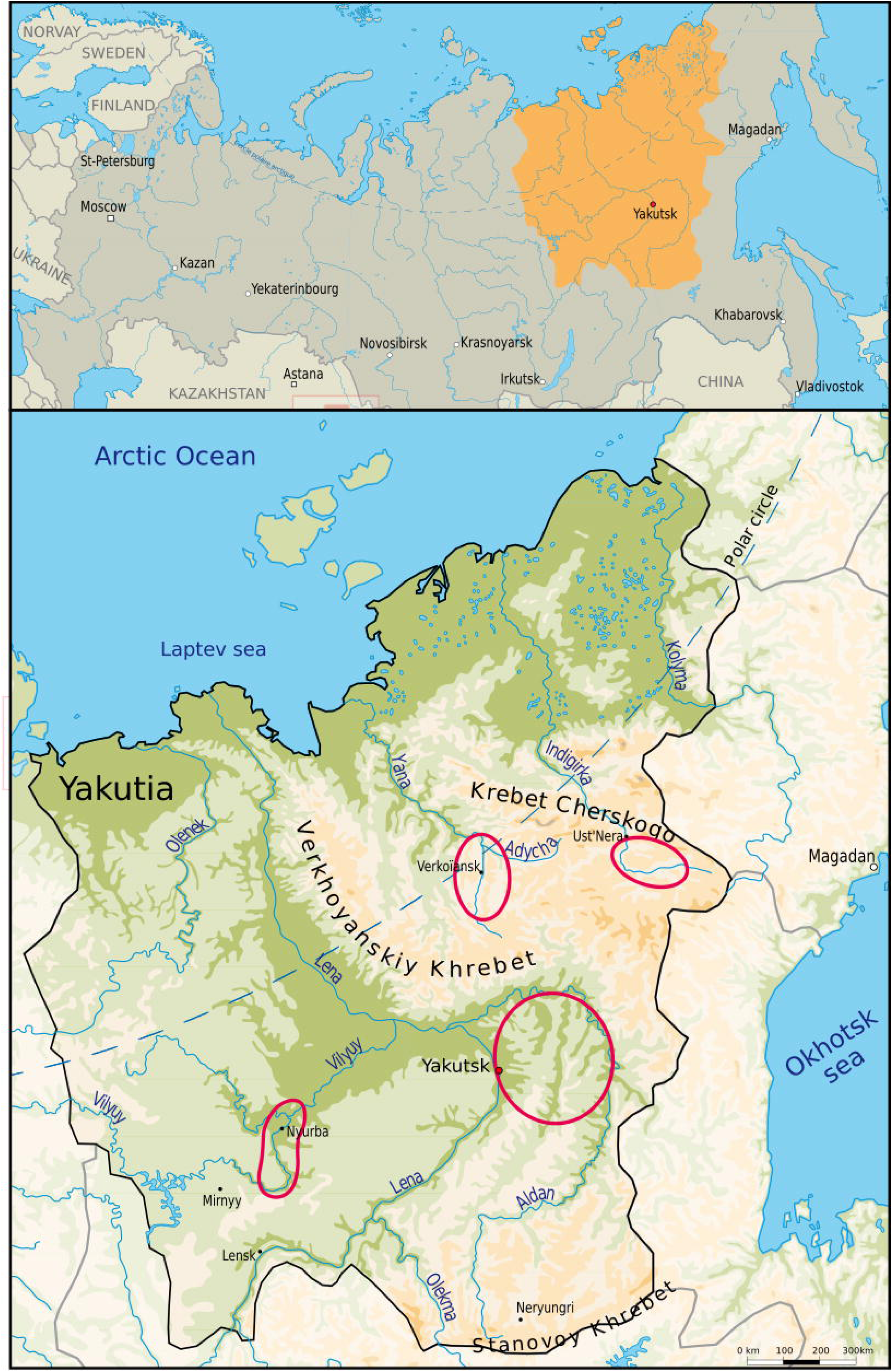

## Materials and methods

### Archaeological material

We analysed all clothes and archaeological artefacts present in the graves, in order to compare cultural transmission patterns and population genetic structure using the distribution of Y-haplotypes, Y sub-haplotypes and related pairs of individuals. Factorial correspondence analysis and Ascending Hierarchical Classification (AHC), using Euclidian distances and aggregated around the barycentre of each class (XLSTAT 2016.07.39157), were conducted on the entire sample and the process was repeated on a sample restricted to the 45 adult males for whom full archaeological data, including clothes, was available. 74 parameters were defined: the location (4 regions), dating (4 chronological phases), biological data (8 Y-chromosomal haplogroups), the type and number of artefacts and their modifications according to funerary rituals (42 items), the quality of clothes and the presence of adornment (16 items). This information is presented in Supplementary data II.1.

### DNA typing and sequencing

In previous studies, all ancient individuals and 134 modern individuals were analysed at 15 autosomal STR (Short Tandem Repeat) loci with the AmpFLSTR1 Identifiler Plus kit (Life Technologies™). A total of 56 modern individuals from settlements along the Indigirka were analysed using the Qiagen Investigator 24plex QS1 kit at 21 autosomal STR loci, (D8S1179, D7S820, D3S1358, D13S317, D16S539, D2S1338, D19S433, D5S818, D21S11, CSF1PO, vWA, THO1, TPOX, D18S51, FGA, D1S1656, D12S391, D2S44, D10S1248, D22S1045 and SE33) including all 15 loci typed by Identifiler, along with three gender identification markers (amelogenin, DYS391 and a Y-indel). All males from the ancient samples and 133 modern male individuals were genotyped at 17 Y-STR loci using the AmpFLSTR1 Yfiler kit (Life Technologies™) (Thèves *et al.*, 2010). The DNA of 34 modern Yakut males (Zvenigorosky *et al.*, 2016) and 31 ancient Yakut males sharing the same 17 Y-chromosomal STR haplotype was analysed at 24 Y-chromosomal STR loci (DYS19, DYS385a/b, DYS389I/II, DYS390, DYS391, DYS392, DYS393, DYS437, DYS438, DYS439, DYS448, DYS456, DYS458, DYS635, Y GATA, DYS449, DYS460, DYS481, DYS518, DYS533, DYS570, DYS627 and DYF387S1a/b) using the AmpFLSTR Y-Filer Plus kit (Life Technologies™). All STR products were run on the 3100 or 3500 genetic analysers (Life Technologies™) and analysed using GeneMapper v. 4.1 (Life Technologies™).

The mitochondrial HV1 region (16024–16383) was amplified in two overlapping fragments and sequenced as described in (Keyser *et al.*, 2003) as a confirmation tool for maternal relationships (Keyser *et al.*, 2015). Haplogroup assignment deduced from the HV1 haplotype from the phylotree system (www.phylotree.org) was confirmed by the typing of SNPs of the mitochondrial coding region. Thirteen SNPs at nucleotide positions 663, 13263, 1715, 5178, 752, 1438, 13086, 7028, 14766, 13708, 4917, 12308, and 9090 were selected, combined in one multiplex PCR reaction and typed using the iPLEX1 gold technology (Sequenom) as described in (Keyser *et al.*, 2009).

### Genetic kinship testing

Likelihood Ratios (LR) were used to identify all possible familial relationships among ancient samples (Keyser *et al.*, 2015). The markers used were the STR loci previously described, and the allelic frequencies used were calculated using unrelated ancient samples. All Y-STR and mitochondrial HV1 regions were also analysed, to confirm the conclusions based on LR calculations. Some relationships were left unexplained, as expected when studying remote populations with such numbers of markers (Mo *et al.*, 2016; Morimoto *et al.*, 2016; Zvenigorosky *et al.,* 2016). Because classical kinship testing methods are not effective in identifying distantly related duos (Zvenigorosky *et al.*, 2016), we considered genetic proximity to be indicative in cases where several individuals could be compared and included in the genealogy, with second-degree relationships confirming each other. Some lineages were rare or sometimes unique to individuals that were identified as closely related by LR. In cases of second-degree kinship in which some subjects are missing from the genealogies, parental lineages also invalidate some of the possible configurations, leaving a reduced number of hypotheses. The first tests and theoretical discussions of kinship were described (Keyser *et al.*, 2015; Zvenigorosky *et al.*, 2016), but for the first time we considered the totality of the cases, including lineages and second-degree kinship, as well as the associated geographical and cultural data, to evaluate sampling bias.

## Results 1 – Organising individuals according to archaeological material and situation

### Temporal and geographical distribution of the graves

A total of 31,000 km^2^ were surveyed over 15 years in four regions (Central Yakutia between the Lena, Amga and Aldan rivers, the Villuy River Basin, the region of Verkhoyansk and the Upper Indigirka). Three of these locations had already been settled by the Yakuts at the time of the Russian expansion into Eastern Siberia (1620). The latter (Upper Indigirka) was a pocket settlement prior to the year 1700. 174 graves were excavated, 111 of which anterior to 1800. Most were isolated and contained only one body, but 42 subjects belonged to burial grounds of 2 to 7 individuals and some graves were double or triple graves; one grave contained 5 individuals (Supplementary data IV.2). Under the conservative hypothesis of a population of 50,000 in 1700 (Crubézy and Nikolaeva 2017) and if all individuals had been buried, there should exist 200,000 graves for the 18^th^ century alone, with half the occupants having died before the age of 18. Compared to the number of deceased individuals, the number of graves is exceedingly small.

The analysis of the archaeological data, combined with the absolute dating of the graves and the factorial correspondence analysis previously undertaken on the whole sample, confirms that Yakut culture can be divided into several chrono-cultural phases, defined by artefacts, funerary architecture and rituals (Crubézy and Alexeev, 2007, 2012). Tombs anterior to 1689-1700 reveal no Russian influence, even though Russian presence is acknowledged since 1620. This early period is under-represented in the archaeological record outside Central Yakutia, with only a few graves found, restricted to male individuals. The period that extends from 1689-1700 to 1750 represents the “Yakut golden age”, with burials of both males and females becoming more frequent in the four regions. Some graves were opulent and furnished with large amounts of imported goods and pearls, while others were significantly less furnished, despite belonging to members of a social elite, as suggested by the presence of specific items (Crubézy and Alexeev, 2007, 2012). The third period extends from 1750 to 1800 and marks an economic decline with fewer imported artefacts, the beginning of Christianisation, and the appearance of children’s graves (which had been very rare). After 1800, Christianisation reached the core of the population and traditional grave artefacts tended to disappear, with progressive assimilation into graveyards, in a Russian Orthodox fashion (Crubézy and Nikolaeva, 2017). For this last period, a very large number of graveyards with several burials have been identified but only isolated graves outside graveyards have been excavated. Starting in this period, graveyards seem to represent the natural mortality of the population, which had never been the case before in Yakut history.

We separated the 45 adult males for whom complete archaeological data was available into three clusters using Ascending Hierarchical Classification (Figure 4 below). (i) The first group (14 subjects) gathers all graves that presented large numbers of artefacts, mostly (12/14) from the 1700-1750 period. The factorial analysis (Supplementary data II.2) particularly underscores the similarity between those wealthy graves that also yielded signet rings associated to clan chieftains (Crubézy and Nikolaeva 2017). The engravings on these rings are different, although a Central Yakutia subject and a Verkhoyansk subject wore similar rings. Lower hierarchical levels only distinguish between regions. A single grave within the cluster is separated from the others because, although it shares all its cultural particularities with the others, the clothes of the subject buried include far fewer pearls than the others. The two graves anterior to 1700 originate from the same cultural tradition as the 1700-1750 elite and they aggregate with graves from the Verkhoyansk group. (ii) The second cluster (9 subjects) gathers Christian graves from all regions, for which the significant separator was “quality of the ritual”, characterised by the use of candles and crosses. (iii) The third and last cluster (22 subjects) comprises disparate graves from all regions and periods that display associations of diverse artefacts. Most of the graves anterior to 1700 (12/14) belong to this cluster, an indicator of the diversity of grave goods and presentation of the corpse in these graves.

## Results 2 – Organising individuals according to Y-chromosome lineage (male line)

The Y-chromosome of 72ancient males was studied at 17 Y-STR loci and their profiles were compared and organised into haplotypes and groups of haplotypes (Figure 2 – Network). We used the estimated period of their burial to establish possible links between epochs and geographical locations (Supplementary data III.3). For all males (34) that shared the most common profile (Ht1) and the second most common profile (Ht2) we studied 7 rapidly-mutating markers to distinguish mutations between two consecutive generations.

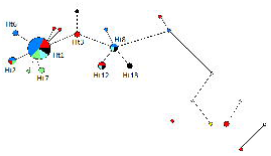

### Y-chromosome diversity

We foundthat most 17-STR profiles only differ from each other by one mutated position, across all epochs and locations (Supplementary data III.1). Furthermore, there is very little diversity and several haplotypes are represented by more than one individual. This observation is only partially explained by the presence of related individuals in the graves that were studied. Many male subjects share a paternal line even though they do not share a first-degree relationship (Keyser et al., 2015).

The main feature of this analysis is that one central haplotype (Ht3) is the most likely ancestor of most other important haplotypes, namely Ht1 and Ht8, which in turn are the most likely ancestors of other peripheral haplotypes. We endeavoured to estimate the time of divergence (using the Network software, Copyright Fluxus Technology Ltd 1999-2017) between the different haplotypes but given the low diversity and the limited timescale (only a few centuries between the most ancient and the most recent subjects), results do not provide precise information. Nevertheless, half (10 out of 20) of all haplotypes, including the most common haplotypes, were already present in the population before the year 1700. This confirms that divergence between the different haplotypes (whichever one being the original haplotype) must have occurred before 1700. This does not inform us as to the place of the divergence, as peripheral haplotypes may have arisen before the carriers had colonised Yakutia (possibly in the 13^th^ century according to archaeological data) or after they had already settled there.

This network study also informs us that 4 haplotypes appear to be separated from the bulk of the population by six to twenty-one (one-step) mutations. These 4 haplotypes belong to 5subjects from the pre-1700 period, which are dispersed on a peripheral branch (MuseeEthno, Byljasik3, OyogosseTumula I, Balyktaek and Cepzeney). These subjects are featured in the relevant figures but they only serve to indicate that some sites yield graves that do not belong to the typical Yakut lineages, although these are sometimes not distinguishable without genetic analysis.

The network study of the sub-haplotypes of Ht1 (i.e. the 24 Y-STR profiles of the subjects that belonged to haplotype Ht1 as previously defined) provides a similar result (Supplementary data III.2). Most sub-haplotypes were already present at the start of the studied period and very few subjects differ from the main sub-haplotype (Ht1s1) by more than one mutation. The maximum number of mutated positions (one-step) is three.

### Geographical and temporal divisions

We studied the geographical repartition of Y-haplotypes (on 17 Y-STR) throughout Yakutia and through time (Supplementary data III.3). It is apparent that the dominant haplotype Ht1 is not confined to one location or time and this is also true of Ht2, Ht8and of Ht12. Three other haplotypes (Ht6, Ht13 and Ht17) are more specifically confined to only two unrelated individuals from one region and one 50-year period. Two haplotypes are represented by unrelated individuals from the same period in different regions (Ht3, Ht16). All other haplotypes are unique.

Ht1, which was found in every region, was divided by the analysis of 7 extra Y-STR markers into eight subhaplotypes (Ht1s1 to Ht1s8, Supplementary data III.4), three of which corresponded to several individuals. Ht1s1 is present in graves across Yakutia, except in the region of Verkhoyansk. Ht1 men found in Verkhoyansk were not Ht1s1 but Ht1s2, a sub-haplotype that was found only in graves from that region and two subjects from the Villuy region. Ht1s2 first appears in the archaeological record in the 18^th^ century and is still found in the modern population. Ht1s3 can be considered a unique sub-haplotype, as it belongs to three brothers found in adjacent graves (the Shamanic Tree family).

Ht2 is divided into two sub-haplotypes, Ht2s1 and Ht2s2. Ht2s1 belongs to a father and his son who lived around the beginning of the 18^th^ century in the region of Verkhoyansk. Ht2s2 belongs to one of their Verkhoyansk relatives and two subjects from Central Yakutia whoalso lived around the beginning of the 18 ^th^ century.

### Modern data

Among the 146 unrelated Yakut men who took part in the studies over the years, 33 carried the Ht1 haplotype. This represents a proportion of 22.6%, while the proportion of Ht1 men in archaeological data (over all periods), was 47.2% (34/72). Despite important variations, Ht1 remains the most common haplotype in the Yakut population today.

The other multi-regional and multi-period haplotypes were Ht2, Ht8 and Ht12. The first two persisted and are detected in our modern sample, while Ht12 has disappeared. Ht2 declined from 6.9% (5/72) to 0.7% (1/146). Conversely, Ht8 doubled in proportion, from 6.9% (4/72) to 14.4% (21/146).

The haplotypes that were confined to one period and one region (Ht6, Ht13 and Ht17) are all absent from the modern sample. Of the multi-regional, one-period haplotypes, Ht3 survived into modern times and Ht16 did not. Most of the haplotypes that constituted the ancient sample (16/20) did not persist. The four exceptions are Ht1, which was always dominant (with some variation between periods), Ht2, which was present in a wealthy Verkhoyansk family during the early 18^th^ century, Ht3, which was always marginal in our samples but is central in relation to other Yakut Y-haplotypes, and Ht8, a group from Central Yakutia, which was already present from the early 18^th^ century and still found in the late 19^th^ century.

When modern haplotypes are included in the network (Supplementary data III.1), the overview is more complex but similar to the network of ancient haplotypes. Most modern haplotypes are either identical to ancient haplotypes or only distinguished by one or two one-step mutations. The peripheral branch that included 8 marginal individuals is divided in two peripheral branches that include a variety of modern haplotypes, although none are identical to ancient haplotypes. In short, ancient marginal haplotypes are generally not found in the modern sample but similar haplotypes (that differ by only a few mutations) are found.

Apart from the punctual disappearance and appearance of minor haplotypes and the higher diversity in the modern sample, there is no indication that the qualitative paternal lineage composition of the Yakut population has been dramatically modified between the year 1700 and the 21^st^ century (Supplementary data III.1). The relative proportions have changed significantly between the 17^th^ century and modern times but the progression is gradual, with punctual events of lineage dominance having an impact on the archaeological record. Given the modern population as a proxy for the consecutive ancient populations, Ht1, Ht2 and Ht3 are at times overrepresented while Ht8 is underrepresented during specific periods. Minor haplotypes are very underrepresented in the archaeological record.

Overall, modern and ancient Y-chromosome haplotypes are either identical or closely related and the composition of the modern living male population shows many similarities with the archaeological samples, although they necessarily only provide a partial picture of the ancient Yakut population.

## Results 3 – Organising individuals according to familial links

We studied the familial relationships, some of which had already been discussed in this ancient Yakut sample (Keyser 2015, Zvenigorosky 2016) and focused on their spatial and temporal implications for the history of the population. We focused on the 27 first-degree relationships among the 150 subjects and on the greater number of possible distant relationships, from half-siblings to first or second cousins. The latter could only be confirmed when three or more individuals were involved (e.g. a trio including a mother, her son and her brother provides confirmation for the uncle-nephew relationship).

### Notable features of the families identified

Large families were among the 150 subjects that were excavated and they all originated from two regions, Central Yakutia and Verkhoyansk (Supplementary Data IV.1). Most other genetic links identified concerned individuals that were buried in the same grave or nearby graves and most were also found in the same region ((Supplementary Data IV.2). The four large families (that included more than two individuals) allowed us to make assessments of funeral decisions and to determine the interactions between lineages.

#### Shamanic Tree in Central Yakutia

The most notable family (Figure 3) was made up of five individuals buried in the same grave and a sixth subject buried nearby, in the vicinity of a shamanic tree (ref Keyser) in Central Yakutia. This “Shamanic Tree Family” included an old woman, her daughter and son, and the three infant sons of that daughter. The five subjects were buried simultaneously in the collective grave and the last son died several years later. Kinship testing confirmed the genealogy without ambiguity, except for the status of the last son, who could have been the other children’s half-brother (ref Zvenigorosky). All individuals carried the majority mitochondrial haplotype, transmitted by the oldest woman to all her descendants, and the adult man (the uncle of the children) carried the Ht1s1 Y-chromosome-subhaplotype, which is the main sub-haplotype of Ht1, itself the main haplotype. The children carried sub-haplotype Ht1s3, unique to them and differing from Ht1s1 by a single one-step mutation.

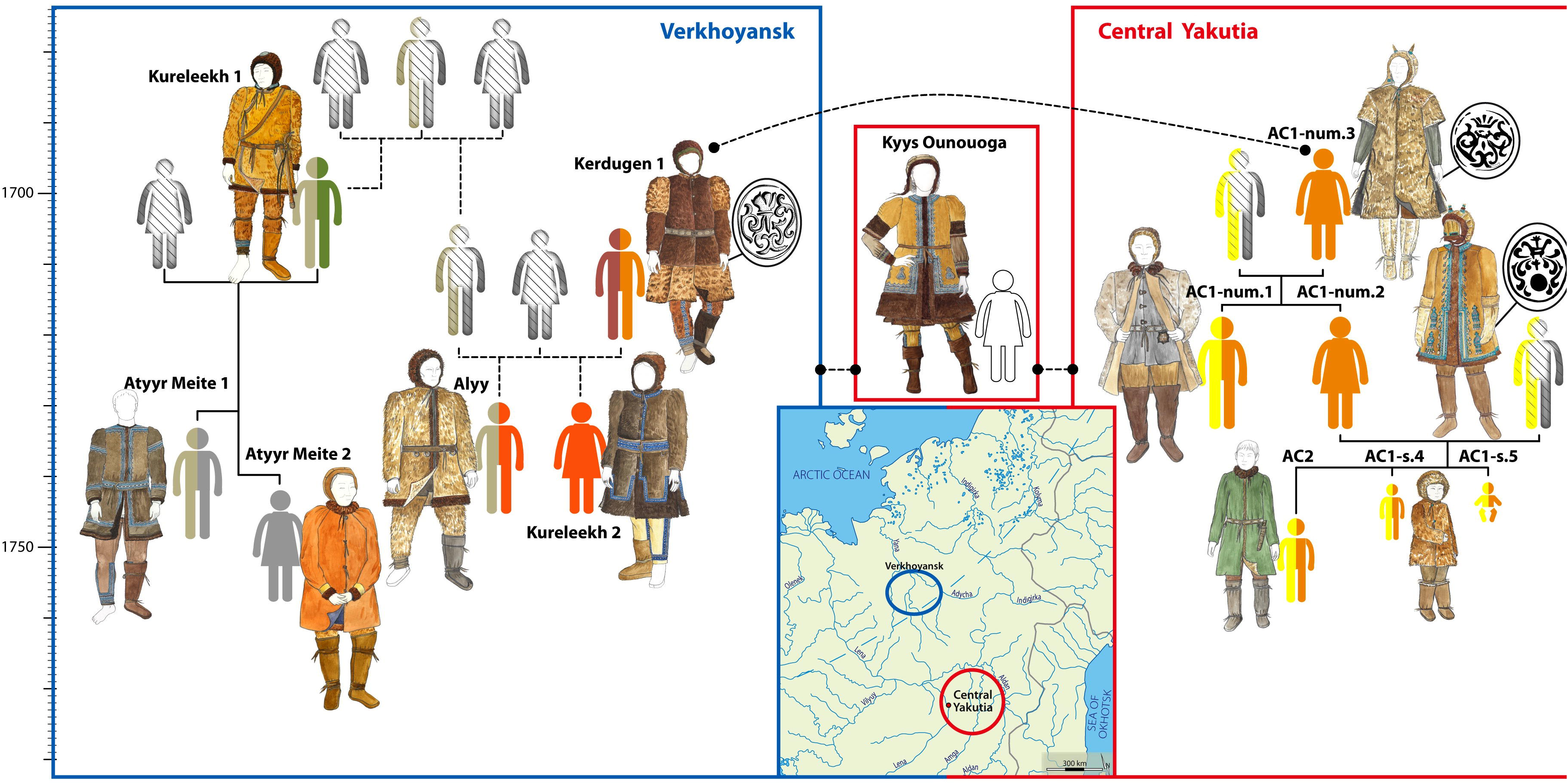

These results indicate that all Shamanic Tree subjects were associated with the majority Y-haplotype Ht1, either because they carried it themselves or because they were related to individuals that carried it, biologically or through marriage.

#### Lepsei and Kerdugen in Verkhoyansk

The region of Verkhoyansk yielded two notable families (Figure 3). The first (“The Lepsei Family”) was composed of a mother and her two new-born sons. The mother probably died in childbirth. The woman’s brother was also buried 300 metres away. He carried the majority haplotype and sub-haplotype Ht1s1, while the children carried the(now extinct) haplotype Ht7. The second family (“The Kerdugen Family”) presented a more complex case. It included six individuals, grouped in two closely related trios with uncertain ties to one another. Three graves were found containing a man, his son and his daughter (320 metres apart) and three other graves included a man (Kerdugen 1), his daughter (Kureleekh 2) and a man closely related to her (Alyy). Kinship testing indicated a sibling relationship between Kureleekh 2 and Alyy but they did not share the same mitochondrial DNA. The man Kerdugen 1 was also closely related to Alyy but they did not share either a male line or a female line. The specifics of the case remain unresolved but three out of the four men included in this genealogy carried haplotype Ht2, which is still extant today, although it was only found in one modern individual.

#### The At-Daban/Myran Family in Central Yakutia and Verkhoyansk (supplementary data IV.3)

Two trios, in Central Yakutia and Verkhoyansk, were found to be related to each other. The first was made up of the woman At Daban 0, her son SytyganeSihé 1 and a third man SytyganeSihé 2, who was found to be related to both subjects, although the nature of that relationship was not clear. Both men carried the Y-sub-haplotype Ht1s1, the majority sub-haplotype. The mother-son pair carried the majority mitochondrial haplotype and SytyganeSihé 2 carried a minority mitochondrial haplotype. A second trio uncovered in Verkhoyansk included a mother and her son (Myran and Kouranakh), as well as a third (female) subject, Tysarastaakh 1. All three carried the same mitochondrial haplotype, which was not found in the Yakuts outside of that family. The relationship between Tysarastaakh 1 and the other individuals is not clear, although kinship tests excluded a first-degree (parental or sibling) relationship, while also confirming that she was related to them.

The two trios (the family of At-Daban and the family of Myran) were determined by kinship tests to be related through the relationship between At Daban 0, SytyganeSyhé 1 and Myran. Because tests indicated a probable second-degree (uncle-niece, half-sibling, grandparent/grandchild) relationship between the three, the most parsimonious genealogy presents At Daban 0 as the grandmother of Myran, and SytyganeSihé 1 as her uncle.

### Regional clans and families

The Y-chromosome lineages that are carried by many individuals in our ancient sample mostly issue from families, with two exceptions: Ht1 and Ht1s1, which are very common in the archaeological record and the modern sample, and Ht8, the exhumed carriers of which do not seem to share identifiable parental relationships.

Some lineages appear to share familial links spanning distant regions. The two Central Yakutian families (“Shamanic Tree” and “At Daban”) carry majority haplotypes both on the male and the female line. All men are Ht1 and all women carry the main mitochondrial haplotype. Both families also include adult Ht1s1 men. They appear to be linked to subjects carrying the Ht1s2sub-haplotype in Verkhoyansk through second-degree relationships. Another grave, “KyysOunouoga” (Crubézy and Alexeev, 2007,2012), held an individual that appeared to be related to both the Ht1s1 individual from Shamanic Tree and one Ht2 individual from the extended Kerdugen family. There again, kinship tests did not provide a precise estimation of the link but eliminated close ties and the absence of ties. Because Ht2 is itself linked to Ht6, the regions of Central Yakutia and Verkhoyansk are connected by genealogy, with the ever-dominant Ht1s1 line from Central Yakutia having distant links with three lineages from Verkhoyansk: Ht2, Ht6 and later Ht1s2.

## Discussion

The combined analysis of cultural and geographical characteristics (Figure 4, Hierarchical analysis), along with the consideration of paternal lineages and familial links, indicates that the first cluster of culturally similar ancient Yakut men (from the 1700-1750 period) is composed of 14 subjects carrying 4 Y-chromosome lines. The majority line Ht1 (8 subjects) is found in Central Yakutia and in the Villuy river basin, while Ht8 is found on the periphery of Central Yakutia, in a territory historically peopled by a distinct tribe (the region and tribe of Tataa). Ht2 and Ht6 were found in the region of Verkhoyansk, in two different river valleys. The study of the DNA of the subjects that could not be included in the archaeological analysis reveals that 4 men from Verkhoyansk carried Y-haplotype Ht1 but they all belonged to a specific sub-haplotype (i.e. using 24 Y-STR), Ht1s2. Therefore, it seems that, in the early 18^th^ century, there existed a differentiation between regional clans, especially in Verkhoyansk. Territories defined by watersheds may be at the origin of this separation of Y-chromosomal haplotypes. Among Ht1 men, only Ht1s1 individuals wore chieftain rings with engravings. The only other man to have worn a chieftain ring was Kerdugen, of the Ht6 line in Verkhoyansk. The engravings on his signet ring are similar to those found on the signet ring of one of the relatives of the Shamanic Tree family in Central Yakutia. There were other familial relationships between Ht1s1 subjects in Central Yakutia, who wore signet rings, and the clan chieftains found in wealthy graves in Verkhoyansk, who belonged to Y-chromosome lines Ht2 and Ht6 (Kerdugen 1 and Kureleekh 1, see Figure 3).Only KousTcharbyt (of the Ht8 line on the periphery of Central Yakutia) showed no genetic links with Ht1s1 subjects. He manifestly belonged to a less important elite. In the 18^th^ century, we observe the preponderance of two clans from the Ht1 haplotype group (namely Ht1s1 and Ht1s2), with both men, who carry the haplotype, and women, some of whom are the mothers or daughters of Ht1 individuals and therefore the actors or the fruit of matrimonial alliances. In historical terms, the beginning of the 18^th^ century corresponds to a partial expansion and domination of the Meginzy tribe, whose territory was part of Central Yakutia (Crubézy and Nikolaeva 2017). We endeavoured to identify a possible association of the Ht1s1 haplotype and the Meginzy clan. The woman At Daban 6 was buried in the necropolis of the elite of the Meginzy clan with items, especially rings, which could have been offered to her husband, M. Bozekov, head of the Meginzy clan, by the Tsar or from a noble family in Russia, 6000km away (Supplementary Data IV.4). Her son, SS1, carries sub-haplotype Ht1s1, which indicates that his father also carried this sub-haplotype. Furthermore, polygyny was frequent in the Yakut elite in the 18^th^ century and could easily produce a situation where one grandfather could live alongside as many as 70 sons and grandsons, all sharing his Y-chromosomal profile (Tokarev, 1945). These factors explain how a particular lineage could heavily dominate the archaeological record. Some Central Yakutia individuals, however, seem to have migrated between the two regions. Myran 1, the probable granddaughter of At Daban 6, was buried with her son in Verkhoyansk, 640 km from Central Yakutia. It appears that migrants acculturated with time, rather than replaced local traditions with their own. Subject Kureleekh 1 is a possible example of an original settler in Verkhoyansk: he is the only subject in Verkhoyansk who was buried with a blanket of birch bark, traditional in Central Yakutia but absent in other Verkhoyansk graves, birch being absent from the environment. His clothes and grave artefacts are typical of Central Yakutia and he was buried in a landscape also typical of Central Yakutia but found only rarely in Verkhoyansk. His son AtyyrMeite 1 died at a young age and was buried without a blanket of birch bark, not long after his father, 64 km away on a terrace overlooking a river; an unusual burial place. He had typical Central Yakut clothes, except for reindeer skin trousers, which are typical of the Verkhoyansk region, where reindeer are common. The items in his grave are comparable to his father’s but their disposition is unique, showing that some traditions were disappearing. Kureleekh’s daughter, AtyyrMeite 2, died as an old woman and was buried a few hundred meters away from her brother, in a reindeer skin coat and a grave constructed in a fashion typical of Verkhoyansk. Overall, these are signs of practices changing with the environment and adaptation to new local traditions. However, this does not mean that genetic links between those distant regions were completely severed, as is shown by the case of the families of Shamanic Tree and Kerdugen-Kureleekh.

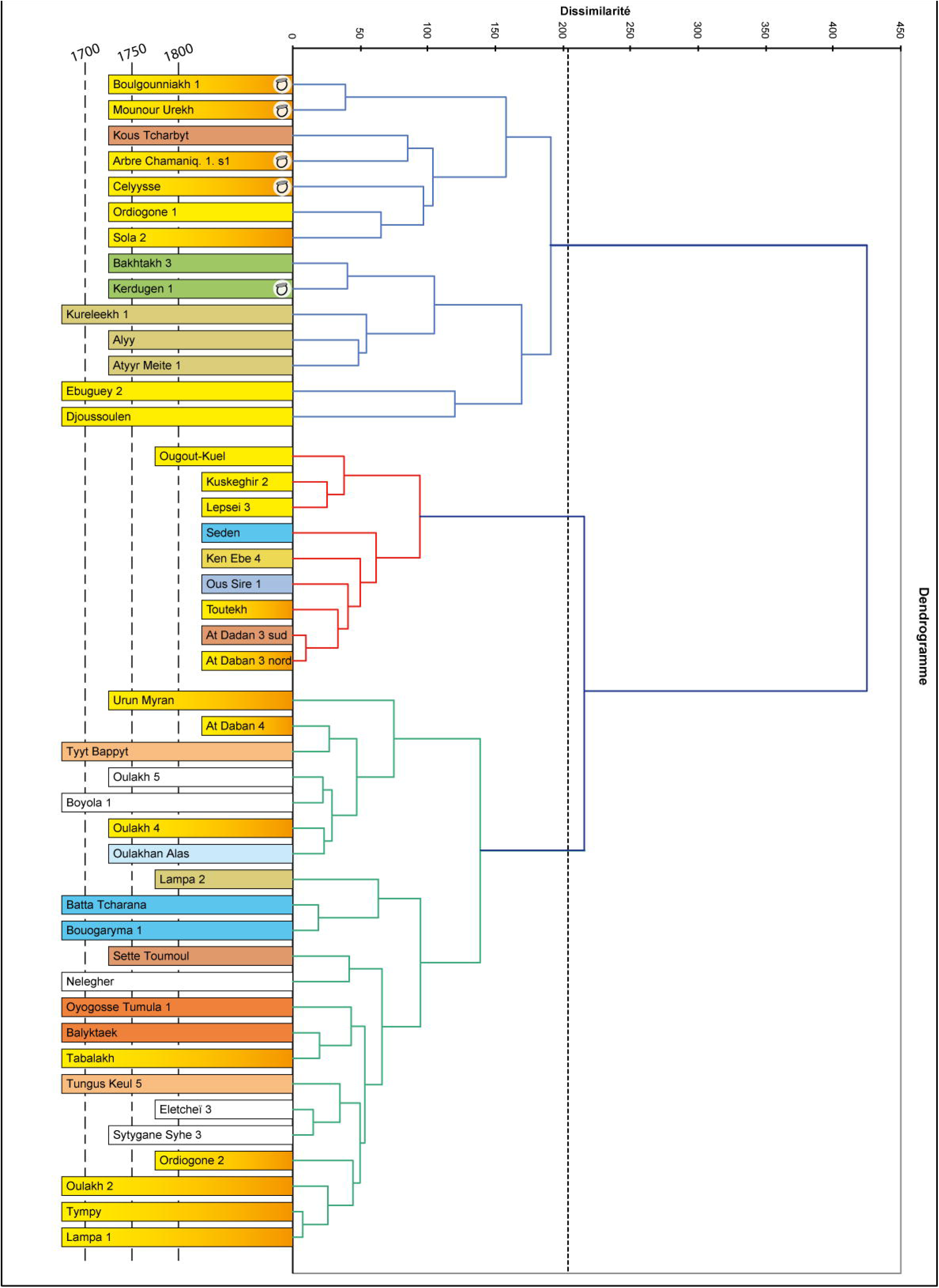

The second cluster, that aggregates Christian graves from all regions, comprises 5 Ht1 men (out of 9), including the most ancient Christianised subject. This suggests that men from the Ht1 line were among the first to be Christianised by the popes looking for elite individuals. It also appears that these individuals continued to expose their wealth in death, even after religious conversion. Christianisation was a force for change, with socio-cultural transmission from missionaries often starting with a single selected individual. In Verkhoyansk, the grave of Christian woman Myran 1, who was married to a member of the local dominant clan Ht1s2, served as the foundation for the community’s Christian cemetery. Her son died before her and was buried 2 kilometres away, in the traditional Yakut manner and with a mirror on his chest, hidden under his clothes, clearly identifying him as a shaman. Verkhoyansk also yielded the grave of early Christian woman AtyyrMeite 1, whose father Kureleekh 1 was a member of the then dominant clan Ht2s1, before the domination of Ht1s2. She was buried on a watershed separating different tribes and her brother AtyyrMeite 2, who died before her, was buried 310m away, following a Yakut ritual. If our comparison had been limited to the artefacts or the mode of burial, these subjects would have been classified as members of different cultural groups and it would not have been suggested that they were closely related and even less part of the same generation. Missionaries may have approached Christianisation through the conversion of elite individuals, possibly preferentially women. That the son of Myran 1 had been a respected shaman could have facilitated the founding of the community cemetery: the conversion of the mother of a shaman from a dominant clan was a success for the popes, which would have made Myran 1 a highly respected Christian. We also noticed a similar rapid change in one generation in the case of the Shamanic Tree family, who was decimated by a smallpox epidemic(Biagini et al.,2012).The third (surviving) child, who was buried 10 years after his mother, was interred without offerings, as in Christian rituals, and his clothes were made of imported fabrics.

The third cluster is very heterogeneous in terms of archaeology and Y-chromosome haplotypes. The geographical repartition of the subjects from this cluster and others (who died before 1700 and whose Y-chromosome was analysed but were not included in the archaeological study)evokes the Yakut melting-pot of that time, linked to the co-existence of several tribes in territories that were then separated (Supplementary data V).These diverse Y-haplotypes can be separated in two groups: (i) A group around Ht3, composed of haplotypes diverging from Ht3 by only 1 to 3 mutations, providing an ensemble that persists in the modern population, with some haplotypes still present and all modern haplotypes closely related to ancient haplotypes. (ii) A group of five subjects separated from the bulk of the male population by 6 to 21 mutations, seemingly corresponding to culturally Yakut or proto-Yakut individuals of very different origins. Some (MuseeEthno and Byljasik3) carry a haplotype with no corresponding modern carrier or even carriers of a similar haplotype. This suggests a cultural rupture between the ancient horse-breeders and modern Yakuts. MuseeEthno is a grave which has been found in a settlement of the “little houses” culture, whose links to the Yakut culture were discussed up until now (Gogolev, 1993). Byljasik3) is the most ancient subject found in the region of Verkhoyansk and both local (current) traditions and unique cranial deformations suggest a Central Asian origin for this subject. Other marginal subjects carry haplotypes that are extant in the modern population, or haplotypes similar to them. This is the case for Cepzeney, a subject buried on his side (one of two such cases) in the 16^th^ century and Balyktaek, a 15^th^ century subject from Central Yakutia showing funeral practices reminiscent of the southern regions and OyogosseTumula 1 (the most ancient subject found in the Villuy region, dated to the 17^th^ century). The Yakut melting-pot appeared to have been followed in the early 18^th^ century by a diminution of male lineage diversity, under the influence of the Meginzy tribe. Two pre-1700 Ht1 subjects aggregate with the first cluster of wealthy graves, showing continuity in the Ht1 line. The cultural tradition of the Yakut golden age in the first part of the 18^th^ century is therefore associated with the Ht1 line. It should be noted that the third cluster of miscellaneous graves also comprises 6 subjects of the Ht1 line and the early 18^th^ century. 3 subjects represent persistent ancient traditions and the 3 remaining subjects were buried on the edge of the Meginzy clan necropolis, which suggests that not all Ht1 subjects were socially equal.

## Conclusion

A combination of palaeogenealogy and the study of cultural associations is effective in the characterization of archaeological samples. It allows the identification of cultural traditions and practices in subsets of the population. Population movements within families and clans, as well as alliance strategies, can be better appreciated. It allows us to distinguish between different mechanisms of cultural transmission (Guglielmino 1995). In our sample, vertical transmission within family lines is proving to be a very conservative process (represented by cluster 1). Horizontal transmission among unrelated individuals (individuals that died before 1700 from cluster 3) is showing to be a more random process. This combined approach also highlights cultural transmission from one subject to its many contemporaries (Christianisation and cluster 2). Cultural transmission of traits can also be directly influenced by survival needs in different environments. Finally, these results highlight the limited size of the archaeological sample, which is intelligible through the lens of history. Before Christianisation, most corpses were not buried but were instead laid out on trestles or platforms in trees, structures which left no archaeological trace (Novgorodov 1955, Alexeev 1996).For the 18^th^ century alone, the remains of at least 200,000 individuals were not preserved. There is an important quantitative bias in our study, which is associated to a burial selection bias for gender (only males were buried before 1700), social origin, as well as family and clan preference between 1700 and 1850.

The Yakut study demonstrates the impossibility of a quantitative approach to an ancient dataset constructed from a funerary assemblage, both in archaeological terms and in genetic terms. On the other hand, the relative homogeneity of the ancient Yakut dataset available, which prohibits large-scale population genetics conclusions, is punctuated by qualitative signs of cultural transmission at all levels: alliance strategies, clan dominance and intragenerational changes. These punctual events allow us to reconstruct, in part, the biological and cultural history of an ancient population, despite the missing elements that either have deliberately been discarded, or remain to be discovered.

## Acknowledgments

Administrative and research work were supported by the program of the France-Russia Associated International Laboratory (LIA COSIE number 1029), associating the North-Eastern Federal University (Yakutsk, Sakha Republic), the State Medical University of Krasnoyarsk, the Russia Foundation for Fundamental Research (Moscow, Russia), the University of Paul Sabatier Toulouse III, the University of Strasbourg I (France) the National Centre of Scientific Research (Paris, France) and the Initiative ďExcellence Chaires ďattractivité, Université de Toulouse (OURASI), attributed to L. Orlando. Funding for excavations was provided by the French Archaeological Mission in Oriental Siberia (Ministry of Foreign and European Affairs, France), the North-Eastern Federal University (Yakutsk, Sakha Republic), and the French Polar Institute Paul Emile Victor.

